# Galanin neurons in the hypothalamus link sleep homeostasis, body temperature and actions of the α2 adrenergic agonist dexmedetomidine

**DOI:** 10.1101/565747

**Authors:** Ying Ma, Giulia Miracca, Xiao Yu, Edward C. Harding, Andawei Miao, Raquel Yustos, Alexei L. Vyssotski, Nicholas P. Franks, William Wisden

## Abstract

Sleep deprivation induces a characteristic rebound in NREM sleep accompanied by an immediate increase in the power of delta (0.5 - 4 Hz) oscillations, proportional to the prior time awake. To test the idea that galanin neurons in the mouse lateral preoptic hypothalamus (LPO) regulate this sleep homeostasis, they were selectively genetically ablated. The baseline sleep architecture of *LPO*-Δ*Gal* mice became heavily fragmented, their average core body temperature permanently increased (by about 2°C) and the diurnal variations in body temperature across the sleep-wake cycle also markedly increased. Additionally, *LPO*-Δ*Gal* mice showed a striking spike in body temperature and increase in wakefulness at a time (ZT24) when control mice were experiencing the opposite - a decrease in body temperature and becoming maximally sleepy (start of “lights on”). After sleep deprivation sleep homeostasis was largely abolished in *LPO*-Δ*Gal* mice: the characteristic increase in the delta power of NREM sleep following sleep deprivation was absent, suggesting that LPO galanin neurons track the time spent awake. Moreover, the amount of recovery sleep was substantially reduced over the following hours. We also found that the α2 adrenergic agonist dexmedetomidine, used for long-term sedation during intensive care, requires LPO galanin neurons to induce both the NREM-like state with increased delta power and the reduction in body temperature, characteristic features of this drug. This suggests that dexmedetomidine over-activates the natural sleep homeostasis pathway via galanin neurons. Collectively, the results emphasize that NREM sleep and the concurrent reduction in body temperature are entwined at the circuit level.

**Significance:** Catching up on lost sleep (sleep homeostasis) is a common phenomenon in mammals, but there is no circuit explanation for how this occurs. We have discovered that galanin neurons in the hypothalamus are essential for sleep homeostasis as well as for the control of body temperature. This is the first time that a neuronal cell type has been identified that underlies sleep homeostasis. Moreover, we show that activation of these galanin neurons are also essential for the actions of the α2 adrenergic agonist dexmedetomidine, which induces both hypothermia together with powerful delta oscillations resembling NREM sleep. Thus, sleep homeostasis, temperature control and sedation by α2 adrenergic agonists can all be linked at the circuit level by hypothalamic galanin neurons.

## Introduction

It has been proposed that sleep aids metabolite clearance (1), synaptic down-scaling (2), stress reduction (3) and protection of the heart (4). Disruption of sleep causes many changes in brain gene expression and the blood plasma metabolome (5–7). Perhaps reflecting the fundamental restorative purpose(s) of sleep, the urge to sleep, known as the homeostatic drive, increases with the time spent awake and dissipates during sleep (8). Sleep deprivation causes a characteristic rebound in NREM sleep accompanied by an immediate increase in the power of delta (0.5 - 4 Hz) oscillations (deeper sleep) and amount of subsequent NREM sleep, proportional to the previous time spent awake (8-11). Widely expressed genes (*e.g. Sik3, Adora1, clock, mGluR5, per3, reverbα*) have been found to modulate sleep homeostasis (9, 11–17). Astrocytes and skeletal muscle release messengers which can modulate the process (18, 19). But in mammals little is known about how sleep homeostasis might work at the neuronal circuit level, or even whether the homeostatic drive is primarily locally or globally determined (20–22).

There is strong evidence that points towards the preoptic (PO) hypothalamus as playing a pivotal role (23). During sleep deprivation and recovery sleep, neurons in this area, as well as in the neighboring bed nuclei of the stria terminalis, become excited (24–27). The preoptic hypothalamus also houses circuitry that regulates body temperature (28–30). cFOS-dependent activity-tagging revealed that after sleep deprivation, reactivating the tagged neurons in the preoptic area induced both NREM sleep and body cooling (26). Indeed, NREM sleep induction and core body cooling are linked by common preoptic circuitry (31), and about 80% of brain cortex temperature variance correlates with sleep-wake states (32). On entering NREM sleep, the neocortex of rats and mice cools rapidly (33, 34).

NREM sleep and low body temperature can also be brought together pharmacologically: α2 adrenergic agonists induce an arousable NREM sleep-like profile (26, 35–38), which in humans resembles stage 2/3 NREM sleep (39–41), but with the complication of sustained hypothermia (26, 35). The metabotropic α2A receptor mediates both the NREM sleep-like state and the hypothermic effects of dexmedetomidine (42, 43).

These α2 adrenergic agonists are increasingly favored over benzodiazepines for long-term sedation (44). Although it used to be thought that dexmedetomidine induces sedation by inhibiting noradrenaline release from neurons in the locus ceruleus (43, 45, 46), there is building evidence that this is not the case (26, 36, 47). Dexmedetomidine induces cFOS expression in the preoptic hypothalamic nuclei (26, 48), and can induce sedation even when noradrenaline release from the locus ceruleus is genetically removed (47). Using c-FOS-based activity-tagging, we found previously that dexmedetomidine requires the LPO hypothalamus to induce both NREM-like sleep and hypothermia and because we obtained similar results following sleep deprivation (see above), we suggested that dexmedetomidine-induced sleep/hypothermia probably involved the same neurons as those activated during sleep deprivation/recovery sleep (26).

Given this potential overlap, the question is whether specific cell types can be identified in the preoptic hypothalamus that are involved in recovery sleep, hypothermia and the actions of α2 adrenergic agonists. Here, we show that the neurons mediating sleep homeostasis after sleep deprivation and dexmedetomidine-induced NREM-like sleep are LPO neurons that express the inhibitory peptide galanin. In mice with selectively lesioned LPO galanin neurons, body temperature is permanently elevated and the sleep-wake cycle is heavily fragmented. Without galanin neurons sleep homeostasis is blunted (no increase in delta power) and the ability of dexmedetomidine to induce high-power NREM-like oscillations and sustained hypothermia is substantially diminsihed. Thus, recovery sleep after sleep deprivation and the deep NREM-like sleep and hypothermia induced by α2a agonists depend on the same hypothalamic circuitry.

## Results

### Selective genetic ablation of mouse lateral preoptic galanin neurons

To selectively ablate LPO^Gal^ neurons, we bilaterally injected a Cre-activatable AAV expressing Caspase 3 (AAV-*FLEX-CASP3*) into the LPO area of *Gal-Cre* mice to generate *LPO*-Δ*Gal* mice (Fig. 1*A*). To confirm LPO^Gal^ neuron ablation, we mixed AAV*-FLEX-GFP* and AAV-*FLEX-CASP3* viruses (Fig. 1*A*). As controls, *Gal-Cre* gene-positive littermates were injected only with AAV*-FLEX-GFP* virus to generate *LPO-Gal-GFP* mice (Fig. 1*A*). The injection coordinates targeted galanin neurons in the LPO (and partially the edge of the MPO area). In the *LPO*-ΔGal mouse group immunohistochemistry with GFP antibodies showed that after five weeks the *AAV-FLEX-CASP3* injections eliminated ~98% of LPO^Gal^ cells, as compared with *LPO-Gal-GFP* littermate controls (Fig. 1*B-D*).

**Fig. 1.**
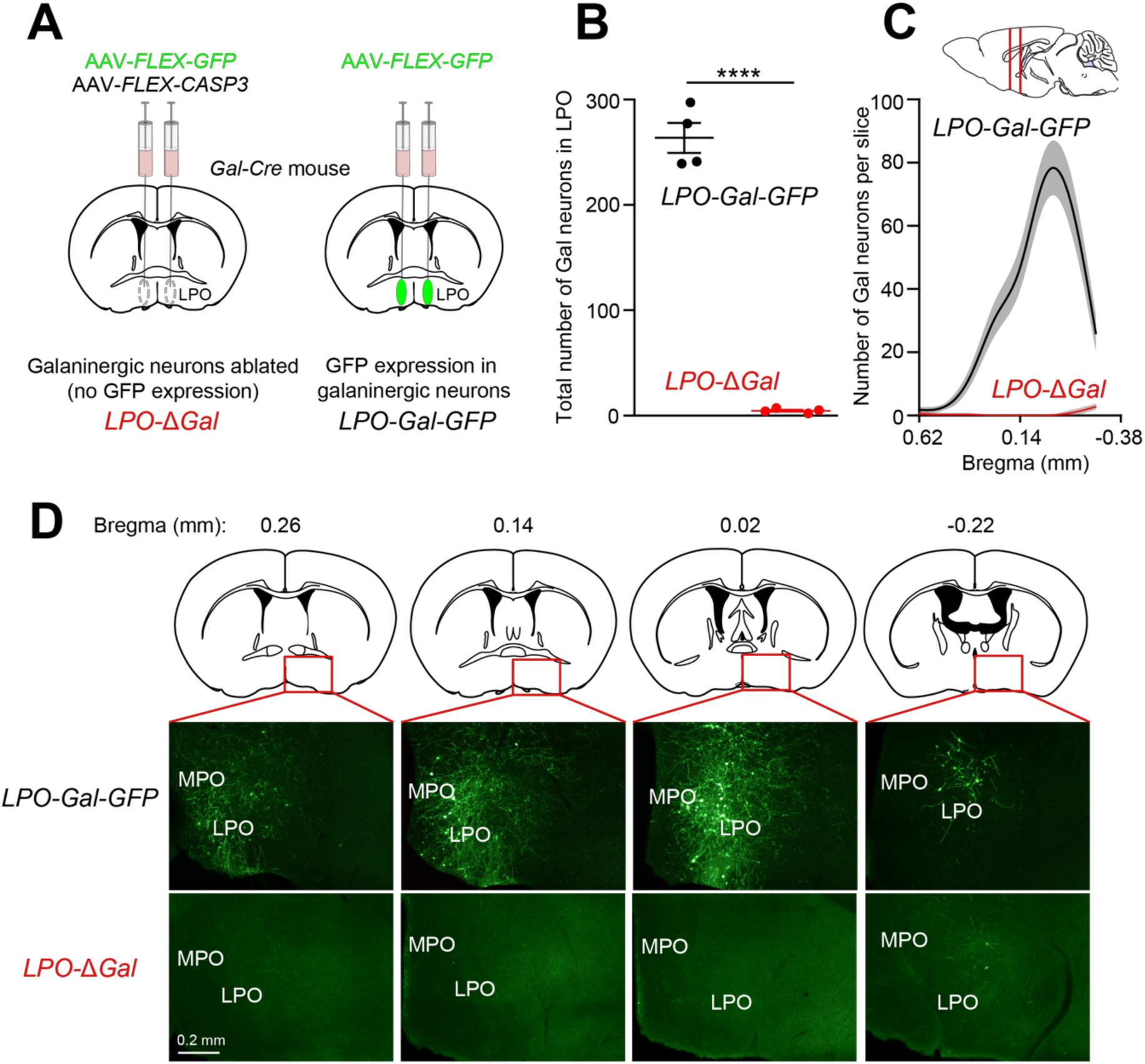
Ablation of galanin neurons in the LPO of the hypothalamus. (*A*) Galanin neurons in the LPO of *Gal-Cre* mice were ablated by bilaterally injecting AAV-*FLEX-CASP3*, together with AAV-*FLEX-GFP* (encoding a fluorescent marker) to give *LPO*-ΔGal mice. Control animals (*LPO-Gal-GFP* mice) were injected with AAV-*FLEX-GFP* alone. (*B*) Most (~98%) galanin neurons in the LPO were ablated (\*\*\*\**P*<0.0001; paired two-tailed *t*-test). (*C*) Rostral-caudal distribution of galanin neurons before and after ablation. (*D*) Example images of galanin neurons in the LPO determined by GFP expression (top row) and after ablation (bottom row). For the data in *B* and *C*, brain sections containing the LPO area from 4 individual mice were selected, *i.e.* 4 brain sections for each coordinate. LPO, lateral preoptic; MPO, medial preoptic. All error bars represent the SEM.

### Selective ablation of lateral preoptic galanin neurons induces a chronic increase in body temperature

Five weeks after ablation of galanin neurons in the LPO area, *LPO*-ΔGal mice had a striking increase in their average core body temperatures compared with *LPO-Gal-GFP* mice (Fig. 2*A, B*). In a continuous recording of body temperature over 5 days, the mice still retained a normal diurnal variation of their body temperature with a higher temperature during “lights-off” period (active phase) and lower temperature during “lights-on” period (inactive phase) (Fig. 2*A*, *B*). However, the average body temperature of the *LPO*-ΔGal mice was raised to 37°C, compared with the average 35.5°C of the *LPO-Gal-GFP* controls (Fig. 2*C*). In addition, the range of body temperature change during the 24-hour cycle increased from 1°C to 2°C. In the *LPO-Gal-GFP* control group, the average body temperature during the day and night was around 36°C and 35°C respectively, whereas the *LPO*-ΔGal group had their average day and night body temperatures around 38°C and 36°C respectively (Fig. 2*C*). Thus, LPO^Gal^ neurons must be acting chronically to induce body cooling.

**Fig. 2.**
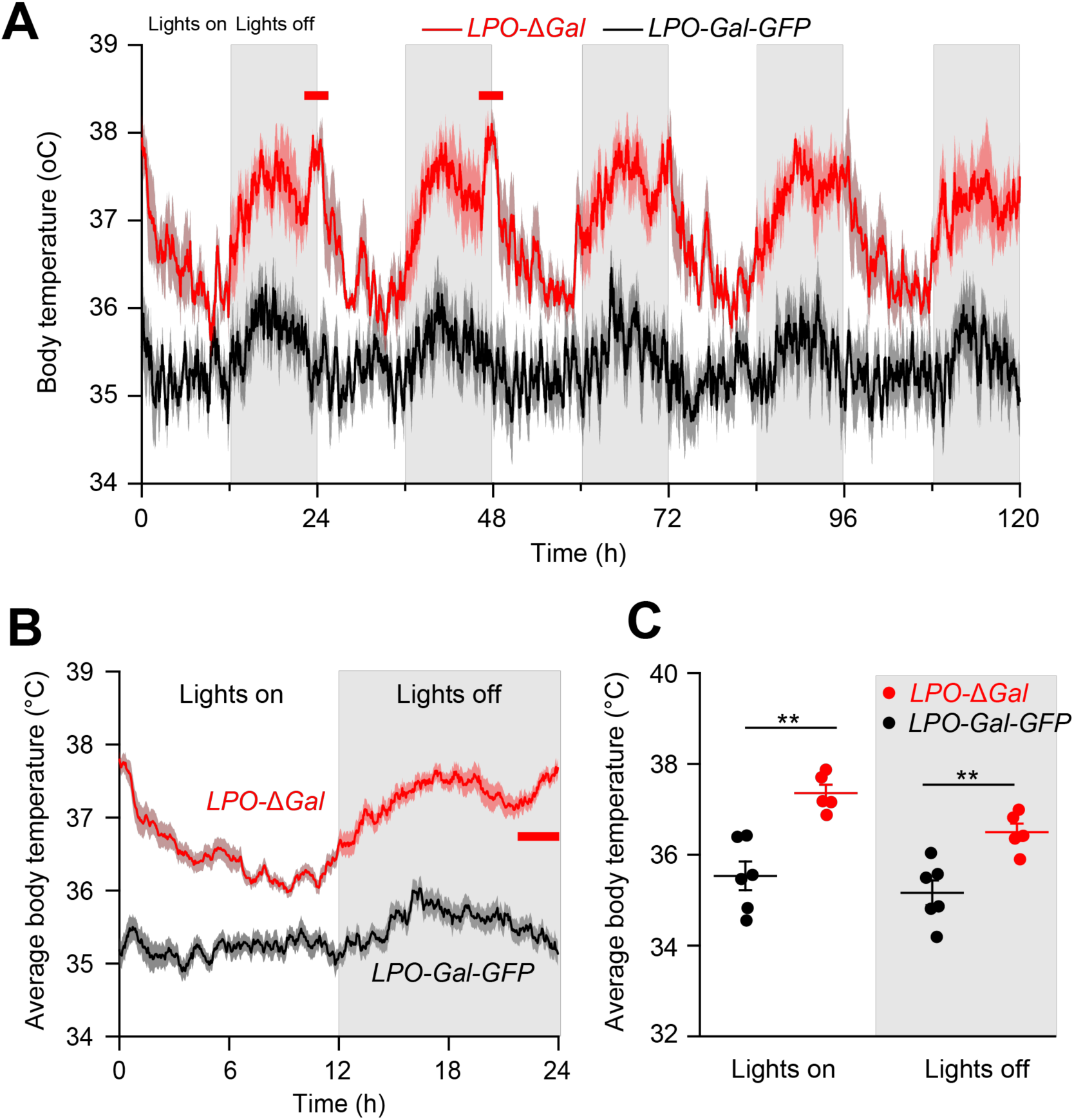
Chronic ablation of galanin LPO neurons markedly elevated core body temperature. (*A*) Ablation of LPO^Gal^ neurons caused increases in both the average core body temperature and its diurnal variation. The record shows typical recordings over five days for both *LPO-Δ*Gal mice (red) and control *LPO-Gal-GFP* mice (black). (*B*) Average core body temperature over 24 hours (*LPO-Δ*Gal mice and control *LPO-Gal-GFP* mice) also shows an abrupt and transient increase in body temperature around the transition from “lights off” to “lights on” in the *LPO-Δ*Gal mice but not the control *LPO-Gal-GFP* mice. (*C*) Average core body temperature increased in *LPO-Δ*Gal mice in both “lights on” and “lights off” (\*\**P*<0.01; unpaired two-tailed *t*-test). (*LPO-GAL-GFP*; *n*=6. *LPO-Δ*Gal; *n*=5). All error bars represent the SEM.

A new feature also emerged in the diurnal temperature variation of the *LPO*-ΔGal mice. In *LPO*-ΔGal mice, a pronounced spike in body temperature appeared just prior to the transition from “lights off” to “lights on”, which was not evident in the *LPO-Gal-GFP* control mice (Fig 2*A, B*, highlighted with red bars) (see Discussion).

Consistent with the above findings, chemogenetic activation of LPO^Gal^ neurons ^with CNO in *LPO-Gal-hM_3_D_q_* mice induced hypothermia (Fig. S1*A, B*), as also reported^ by others (49). CNO had no measurable effect on baseline temperature in control mice (Fig. S1*C*).

### Ablation of galanin neurons in the LPO area increased sleep-wake fragmentation

We examined how LPO^Gal^ neuron ablation influenced the 24-hour sleep-wake cycle (12 hours lights on: 12 hours lights off) (Fig. 3 & Fig. S2). Example EEG and EMG spectra are shown in Figure S2*A*. Chronic ablation of LPO galanin neurons caused a modest reduction in total wake time and an increase in total NREM time during “lights off”, but no change during the “lights on” period (Fig. 3*A*). The amount of REM sleep was unaffected. Furthermore, there were no significant differences in the baseline EEG power in either the WAKE state or NREM state between *LPO-Gal-GFP* and *LPO*-Δ*Gal* mice (Fig. S2*B*). Sleep architecture, however, became highly fragmented following LPO^Gal^ neuron ablation (Fig. 3*B*). The number of WAKE and NREM episodes increased markedly, while their durations were shortened. These effects were most marked during the “lights off” period. The number of REM sleep episodes and their durations were not affected (Fig. 3*B*). The number of WAKE to NREM and NREM to WAKE transitions were significantly increased (Fig. 3*C*), but transitions between other vigilance states did not change.

**Fig. 3.**
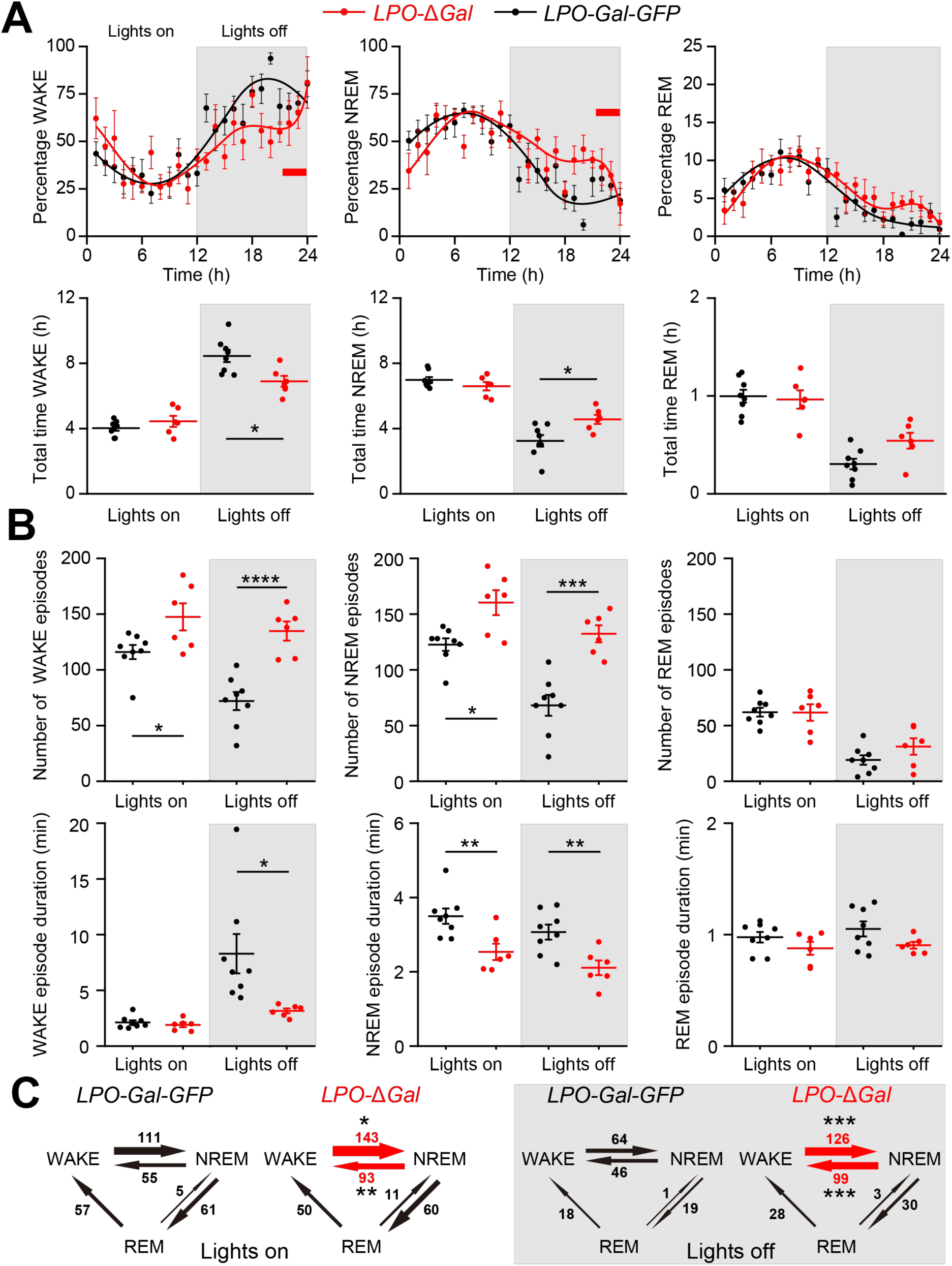
Ablation of galanin neurons in LPO caused profound fragmentation in sleep architecture. (*A*) Ablation of galanin neurons caused only a modest reduction in total WAKE time (\**P*<0.05, unpaired two-tailed *t*-test) and increase in total NREM time (\**P*<0.05, unpaired two-tailed *t*-test) during “lights off”, but no change during “lights on”. The amount of REM sleep was unaffected. (*B*) Sleep architecture, however, was highly fragmented by galanin neuron ablation. The number of WAKE and NREM episodes increased markedly, while their durations were shortened. These effects were most marked during “lights off”. The number of REM episodes and their durations were not affected. (*C*) The number of WAKE to NREM and NREM to WAKE transitions were significantly altered, but transitions between other vigilance states did not change (*LPO-GAL-GFP*; *n*=8. *LPO-Δ*Gal; *n*=6). \**P*<0.05, \*\**P*<0.01, \*\*\**P*<0.001, \*\*\*\**P*<0.0001. All error bars represent the SEM.

Chemogenetic activation of LPO^Gal^ neurons with CNO in *LPO-Gal-hM_3_D_q_* mice induced NREM sleep (Fig. S3 *A,B,C*), as also reported by others (49). The power of this CNO-induced NREM sleep was higher than baseline power of NREM sleep after saline injection (Fig. S3*D*). CNO had no measurable effect on baseline sleep after saline injection in control mice (Fig. S3*E*).

### Ablation of LPO galanin neurons abolishes sleep homeostasis after sleep deprivation

To examine how LPO^Gal^ neurons regulate sleep homeostasis, a 5-hour sleep deprivation was applied to both groups of mice. In control *LPO-Gal-GFP* mice, there was a strong reduction in wakefulness and an increase in total sleep (NREM + REM sleep) following five hours of sleep deprivation (Fig. 4*A*). The main effect, compared to the baseline diurnal variation in wake and sleep times, occurred during the “lights off” period following sleep deprivation (which was carried out during the “lights on” period) (Fig. 4*A*). During the sleep rebound of LPO-Gal-GFP mice, the power in the delta wave band (0.5 - 4 Hz) was also significantly increased compared to the delta power during baseline sleep at the equivalent zeitgeber time (Fig. 4*B, E*). This delta power increase is a characteristic of recovery sleep (8). In *LPO*-Δ*Gal* mice, however, there was no change in WAKE or total sleep (NREM + REM) time following five hours of sleep deprivation (Fig. 4*C*). The increase in delta power seen in control animals following sleep deprivation was also abolished in *LPO*-Δ*Gal* mice (Fig. 4*D, E*). Most (~80%) of the sleep lost as a result of five hours of sleep deprivation was recovered after 19 hours in *LPO-Gal-GFP* mice, whereas only ~22% of the sleep loss was recovered in *LPO*-Δ*Gal* mice (Fig. 4*F*). Indeed, the sleep recovery rate after sleep deprivation was significantly reduced in *LPO*-Δ*Gal* mice compared with *LPO-Gal-GFP* (Fig. 4*F*, *G*).

**Fig. 4.**
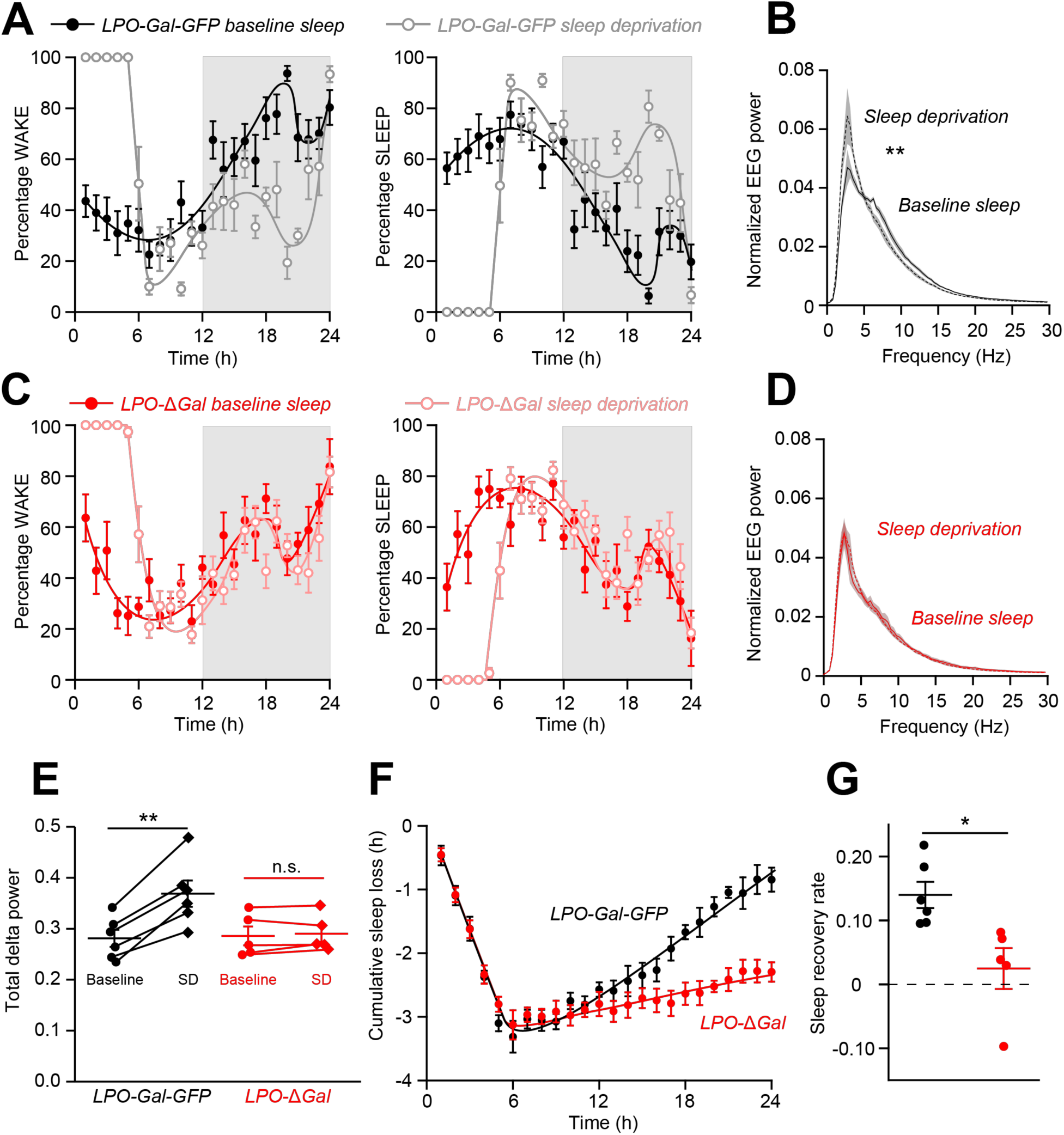
Homeostatic sleep rebound following sleep deprivation was largely abolished by the ablation of LPO galanin neurons. (*A*) In control *LPO-Gal-GFP* mice, there was a strong reduction in WAKE and an increase in total sleep (NREM + REM) following five hours of sleep deprivation. The main effect, compared to the baseline diurnal variation in WAKE and sleep times, occurred during the “lights off” period following sleep deprivation (which was carried out during the “lights on” period). (*B*) During the sleep rebound, the power in the delta wave band (0.5 - 4 Hz) was also significantly increased (\*\**P*<0.01; paired two-tailed *t*-test) compared with the delta power during baseline sleep at the equivalent zeitgeber time. (*C*) In *LPO*-Δ*Gal* mice, however, there was no change in WAKE or total sleep (NREM + REM) time following five hours of sleep deprivation. (*D*) The increase in delta power seen in control animals following sleep deprivation was also abolished in *LPO*-Δ*Gal* mice. (*E*) Quantification of the power spectra in (*B*) and (*D*), i.e. the delta power (0.5 – 4 Hz) after sleep deprivation, showing a significant increase in delta power in *LPO-Gal-GFP* mice after sleep deprivation (P < 0.01, two-tailed paired *t* test, n= 6), but not in *LPO*-Δ*Gal* mice. (*F*) Most (~80%) of the sleep lost as a result of five hours of sleep deprivation was recovered after 19 hours in *LPO-Gal-GFP* mice (black; *n*=6) whereas only ~22% of the sleep loss was recovered in *LPO*-Δ*Gal* mice (red; *n*=5). (*G*) The sleep recovery rate after sleep deprivation is significantly reduced in *LPO*-Δ*Gal* mice (n = 5) compared with *LPO-Gal-GFP (n= 6)* (P < 0.05, two-tailed unpaired t test). All error bars represent the SEM.

### Ablation of LPO^Gal^ neurons strongly reduced dexmedetomidine-induced hypothermia

An unwanted side-effect of dexmedetomidine is that it induces marked hypothermia (26, 35). In control *LPO-Gal-GFP* mice, after injection (*i.p.*) of 50 µg/kg of dexmedetomidine, there was a strong reduction in core body temperature from about 36°C to 25°C over the course of 2 hours (post-dexmedetomidine injection) (Fig. 5*A, B*). This hypothermia persisted beyond 4 hours post-injection. In *LPO*-Δ*Gal* mice, however, the initial reduction in body temperature after dexmedetomidine injection lasted only for the first hour, did not reach the same nadir as in *LPO-Gal-GFP* control mice, and the body temperature nearly returned to baseline levels over the next hour (Fig. 5*A, B*).

**Fig. 5.**
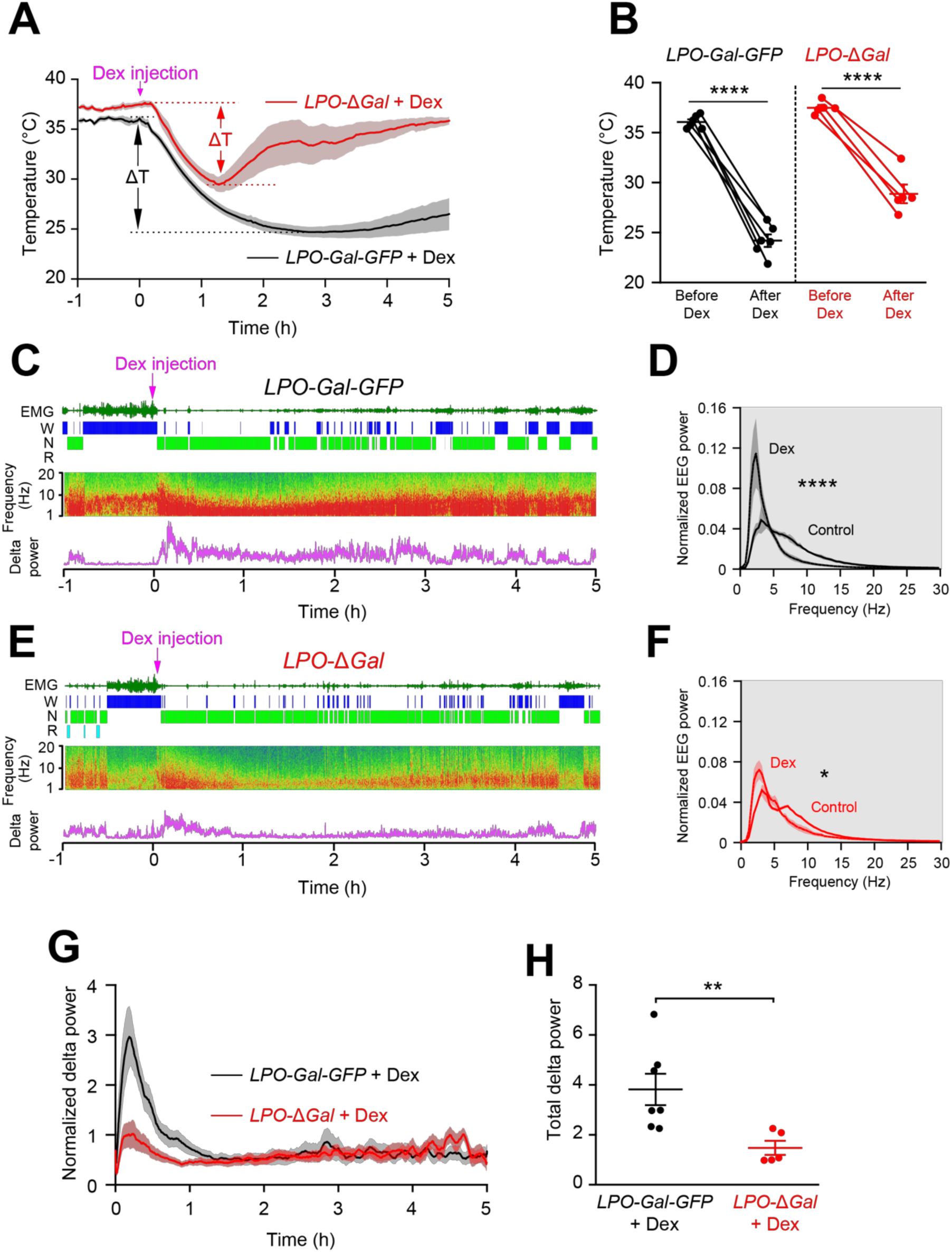
The characteristic sustained hypothermia and increased NREM-like δ power induced by dexmedetomidine were both largely abolished by the ablation of LPO galanin neurons. (*A, B*) In control *LPO-Gal-GFP* mice, there was a strong reduction in temperature from about 36°C to approx. 25°C over the course of 75 minutes (post-dexmedetomidine induction) (*n*=6; \*\*\*\**P*<0.0001, paired two-tailed *t*-test). This hypothermia persisted beyond 4 hours post-injection. In *LPO*-Δ*Gal* mice, however, the initial reduction in body temperature commenced after dexmedetomidine injection lasted only for the first hour, did not reach the same nadir as in *LPO-Gal-GFP* control mice, and the body temperature almost returned to starting baseline levels (33 ± 1.5°C) over the next hour (*n*=5; \*\*\*\**P*<0.0001, paired two-tailed *t*-test). (C, D) Dexmedetomidine injection of *LPO-Gal-GFP* mice: examples of EEG and EMG raw data and vigilance-state scoring and the EEG power spectra (averaged over 30 mins after injection) compared with saline-injected controls. Dexmedetomidine induced a large increase in δ power relative to the NREM sleep baseline (control spectrum). (*n*=6; \*\*\*\**P*<0.0001, paired two-tailed *t*-test). (E, F) Dexmedetomidine injection to *LPO*-Δ*Gal* mice: examples of EEG and EMG raw data and vigilance-state scoring and the EEG power spectra (averaged over 30 mins after injection) compared with saline-injected controls. Dexmedetomidine did not induce a large increase in δ power relative to the NREM sleep baseline (control spectrum) (*n*=5; \**P*<0.05, paired two-tailed *t*-test). (G, H) Time courses of evoked NREM-like delta power following dexmedetomidine *i.p.* administration to *LPO-Gal-GFP* and *LPO*-Δ*Gal* mice. In *LPO-Gal-GFP* control mice, within 20 minutes of injection, dexmedetomidine induced a large increase in delta power relative to the NREM sleep baseline, but this was substantially less in *LPO*-Δ*Gal* mice (\*\**P*<0.01, unpaired two-tailed *t*-test). All error bars represent the SEM.

### Dexmedetomidine requires LPO^Gal^ neurons to induce NREM-like delta power

We concurrently investigated if ablation of LPO^Gal^ neurons compromised dexmedetomidine’s ability to induce a NREM-like sleep state (Fig. 5*C-F*). After one-hour baseline recording, animals received 50 µg/kg dexmedetomidine (*i.p.*) in the “lights-off” period, their active phase. In *LPO-Gal-GFP* control mice, within 20 minutes of injection dexmedetomidine induced a large increase in delta power relative to the NREM sleep baseline (Fig. 5*C, D*), but this increase was substantially weaker in *LPO*-Δ*Gal* mice (Fig. 5*E, F*). Looking at the time course of the evoked delta power following dexmedetomidine injection (Fig. 5*G, H*), in *LPO-Gal-GFP* mice the delta power peaked at 11 minutes post-injection, and then declined over the following hours even though the mice were still in a NREM-like sleep. (Because of the evolving hypothermia in the dexmedetomidine-injected *LPO-Gal-GFP* mice (Fig. 5A), the power of the EEG spectrum declines with time as we and others documented previously; however, the vigilance state can still be scored as NREM sleep-like (31, 50)). By contrast, the delta power remained at baseline NREM sleep levels in dexmedetomidine-injected *LPO*-ΔGal mice, even at the start of the experiment (Fig. 5*G, H*).

## Discussion

Previously, using cFOS-dependent activity-tagging, we found that neurons in the LPO area were sufficient to recapitulate NREM-like sleep and body cooling after both sleep deprivation and dexmedetomidine administration, but we did not identify the cells involved (26). Here, using selective genetic lesioning, we have demonstrated that these are likely to be galanin neurons. Without LPO galanin neurons, the sleep-wake cycle becomes highly fragmented, and sleep homeostasis (the enhanced delta power following sleep deprivation and the extra NREM sleep that follows) is diminished, suggesting that LPO galanin neurons track the time spent awake. Although previously, genes have been identified that modulate sleep homeostasis (see Introduction), we describe here the first neuronal cell type implicated in sleep homeostasis. Throughout the animal kingdom, the homeostatic sleep drive is reflected as changing neuronal activity with the time spent awake (23, 26, 51). Sleep homeostasis at the circuit level in mammals, however, has remained mysterious. It appears to be, at least in part, mediated by extracellular adenosine (9, 14), released by astrocytes (18). Adenosine levels, however, only increase during wakefulness in the basal forebrain and not the PO area (52, 53), and so cannot be the direct trigger for LPO galanin neurons to induce NREM sleep. Skeletal muscle can regulate, by an unknown messenger, sleep homeostasis (19), and could also activate LPO galanin neurons, for example.

We find that galanin neurons in the same LPO area are required for a substantial part of the α2 adrenergic agonist dexmedetomidine’s actions in inducing its characteristic substantially high NREM-like delta power (above that of baseline NREM sleep) and sustained body cooling, characteristics which seem to be an exaggeration of recovery sleep after sleep deprivation. Thus, we suggest that the sleep homeostasis circuitry and the circuitry targeted by adrenergic sedatives are likely to be the same.

The circuitry in the PO area that regulates body temperature seems complex (28, 30). Certain PO neurons (e.g. GABA/galanin-, Glut/NOS1-, PACAP/BDNF-, and TRPM2-expressing cells), respond to immediate external or internal thermal challenge by acutely initiating body cooling or heating (28-31, 49, 54-56), but no information has been available for how genetically-specified PO neurons chronically regulate body temperature. We find that without LPO galanin neurons, the diurnal body temperature rhythms of the *LPO*-Δ*Gal* mice are shifted permanently several degrees higher (Fig. 2A). Thus, galanin neurons are contributing to chronic cooling of the body, correlated with the mice having considerable sleep-wake fragmentation. Lesioning of the rat VLPO area, an extremnely focal lesion, produced chronically less NREM with decreased delta power and decreased REM sleep, but body temperature was unaffected (57). This suggests the extreme ventral part of the PO does not contain temperature regulating cells, at least in the rat.

The so far unexplained active link between body cooling and NREM sleep induction seems tantalizing. It was proposed many years ago that the restorative effects of sleep homeostasis depended on lower body temperature (58). Cooling might be actively linked to sleep because cooling during sleep induces cold-induced gene expression that could remodel synapses or serve some other restorative function (33). An extension of this process would be moving deeper into torpor and hibernation (59), where such gene products are also induced and could be involved in rebuilding synapses on arousal from hibernation (59, 60).

In humans, NREM sleep induction appears when the rate of core body temperature decline is at its maximum (61). A new feature has emerged in the diurnal core body temperature variation we have observed in *LPO*-Δ*Gal* mice: a pronounced positive spike appeared in their body temperature around the transition from “lights off” to “lights on” at ZT24 (Fig 2*A, B*), suggesting that LPO galanin neurons would normally be particularly active in driving down body temperature at the point in the diurnal cycle where sleep pressure is highest at the start of the “lights on” period. This would fit with their role in regulating sleep homeostasis. Increasing sleep pressure during the “lights off” wake period would result in LPO galanin neurons becoming active at this transition to lights on, inducing both sleep and driving down body temperature. This could explain why *LPO*-Δ*Gal* mice are actually more awake compared with *LPO-GAL-GFP* mice at the start of “lights on” (see red bar in Fig. 3A), further emphasizing the link between NREM sleep induction and body cooling.

Conceptually, making LPO galanin neurons selectively sensitive to the excitatory effects of hM_3_D_q_ CNO receptors (Fig. S1 & S3) mimics the actions of dexmedetomidine which can directly excite neurons by Gi-mediated inhibition of hyperpolarization-activated cyclic nucleotide-gated cation channels (62). Both CNO and dexmedetomidine induce NREM-like sedation with enhanced delta power and hypothermia, with the exception that the hM_3_D_q_ receptors are confined to LPO galanin neurons, whereas α2a receptors are widespread. Indeed, the initial phase of body cooling triggered by dexmedetomidine still happens in the *LPO*-Δ*Gal* mice. This is likely because α2A receptors are also found on smooth muscle of peripheral blood vessels and so dexmedetomidine will promote heating loss directly by vasodilation of tail veins. In addition, there will also be central (slower) effects, where cooling is initiated by, for example, activation of galanin neurons via the dorsomedial hypothalamus, rostral raphe pallidus and rostroventral lateral medulla (56), to stop brown fat thermogenesis and induce blood vessel dilation.

For baseline NREM sleep, it is established that GABAergic neurons in the PO area inhibit (for example) the wake-promoting histamine neurons in the posterior hypothalamus to induce NREM sleep (63–65). Based on immunocytochemistry, it was suggested that most rat GABAergic preoptic neurons that project to the histamine neurons in the posterior hypothalamus co-released galanin (66); indeed, galanin directly reduces the firing rate of histamine neurons (67). Activating optogenetically GABAergic (non-galanin) terminals from the PO hypothalamus in the area where the histamine neurons are located induces NREM sleep (64). On the other hand, optogenetic activation of PO galanin neuron soma produced wakefulness (64); this result could be caused by other galanin neuron subtypes (68). Our results on chronic lesioning, are consistent with others who found that acute chemogenetic and optogenetic activation of LPO galanin neurons induces induced both NREM sleep and hypothermia (49). Nevertheless, and even allowing for the high sleep-wake fragmentation that appears in LPO-ΔGal mice, we find that LPO galanin neurons are dispensable for achieving baseline NREM sleep. But given that there are mixed populations of PO GABA (galanin and other peptide-expressing types) that induce NREM sleep (49, 64); and that glutamate/NOS1 cells in MPO/MnPO can induce both NREM sleep and body cooling (31), perhaps this is not surprising

Within the PO hypothalamic area, galanin-expressing neurons have quite different functions: galanin neurons coordinate parental behavior (motor, motivational, social) and mating (69, 70), as well as temperature and sleep (49, 56). We cannot rule out if one type of LPO galanin neuron increases NREM delta power, and another promotes chronic cooling. Intersectional genetics would be needed to further target these cells to resolve this, but this will be a complex challenge. In fact, single-cell profiling and multiplex *in situ* labelling of the PO region found at least seven subtypes of galanin-expressing neuron (68). Most of these subtypes are GABAergic and express the *vgat* gene, but several are glutamatergic because they expressed the *vglut2* gene, and one *vgat*/galanin subtype also expressed tyrosine hydroxylase and the vesicular monoamine transporter (68).

LPO galanin neurons in our study are likely to be GABAergic. We found previously that deletion of the vesicular GABA transporter (vgat) expression in LPO, preventing GABA release from LPO GABAergic neurons, abolished dexmedetomidine’s ability to rapidly induce NREM-like sleep (there was no immediate increase in NREM delta power in the first 10 mins, although after an hour dexmedetomidine could still induce full sleep), suggesting that LPO GABAergic neurons were critical for the initial actions of dexmedetomidine (26). Sustained galanin release would be necessary for the longer-term effects of dexmedetomidine on NREM sleep maintenance and lower body temperature.

In conclusion, based on our lesioning results, LPO galanin neurons are at the intersection of NREM sleep induction and body cooling. Although NREM sleep can still occur without these cells, they are needed for chronically cooling to maintain the normal core body temperature. Furthermore, LPO galanin are needed for sleep homeostasis. A similar result has just appeared for zebrafish, suggesting that sleep homeostasis is a primordial function of PO galanin neurons (71). We propose that the control of sleep homeostasis is actively linked to body cooling. Sustained stimulation of these galanin neurons with the α2 adrenergic agonist dexmedetomidine can induce a slide into a torpor like state (if body temperature is not corrected). Thus, these two processes, sleep homeostasis and α2a receptor sedation/torpor induction could be linked at the circuit level by hypothalamic LPO galanin neurons, which serve, in the initial phase of stimulation, to produce heightened NREM delta power above that of baseline sleep.

## Methods

### Mice

Animal care and experiments were performed under the UK Home Office Animal Procedures Act (1986) and were approved by the Imperial College Ethical Review Committee. *Gal-Cre* mice (Tg(Gal-cre)KI87Gsat/Mmucd) were generated by GENSAT and deposited at the Mutant Mouse Regional Resource Center, stock No. 031060-UCD, The Gene Expression Nervous System Atlas (GENSAT) Project (NINDS Contracts N01NS02331 & HHSN271200723701C to The Rockefeller University, New York) (72). In this mouse line, Cre recombinase expression is driven from a bacterial artificial transgene containing the endogenous galanin gene. All mice used in the experiment were equally mixed genders and had the first surgery at the age of 10-12 weeks. Mice were housed individually. *Ad libitum* food and water were available for all mice and a reversed 12 h:12 h light/dark cycle ("lights on" hours: 17:00-05:00) with constant temperature and humidity.

### AAV transgene plasmids

All AAV transgenes had a flexed reading frame in an inverted orientation, and therefore could only be activated by Cre recombinase. The *pAAV-EF1α-flex-taCasp3-TEVp* transgene plasmid was Addgene plasmid #45580 (a gift from Nirao Shah) (73). The *pAAV-CAG-flex-GFP* transgene construct was Addgene plasmid #28304 (a gift from Edward Boyden). The *pAAV-hSyn-flex-hM_3_D_q_-mCherry* transgene construct was Addgene plasmid #44361 (a gift from Bryan Roth) (74).

### Generation of recombinant AAV particles

All AAV transgenes were packaged in our laboratory into AAV capsids with a mixed serotype 1 & 2 (1:1 ratio of AAV1 and AAV2 capsid proteins) as described previously (75).

### Surgeries and stereotaxic injections of AAV

For surgery, mice were anesthetized with an initiation concentration of 2.5% isoflurane in O_2_ (vol/vol) by inhalation and mounted into a stereotaxic frame (Angle Two, Leica Microsystems, Milton Keynes, Buckinghamshire, UK). Mice were maintained anesthetized on 2% isoflurane during surgery. A heat pad was used during the whole surgery to prevent heat loss. For ablating galanin neurons, the two AAV viruses, *AAV-EF1α-flex-taCasp3-TEVp* and *AAV-CAG-flex-GFP* were mixed in a 1:1 ratio prior to injection while a single virus type was injected for the rest of experiments unless otherwise stated. AAV viruses were delivered using a 10 µL syringe (Hamilton microliter, #701) with a 33-gauge stainless steel needle (point style 3, length 1.5 cm, Hamilton). The injection coordinates (bilateral) for the LPO relative to Bregma were: AP +0.02 mm; ML ±0.75 mm; DV was consecutive starting −5.8 (1/2 volume), −5.6 (1/2 volume). A total volume of 0.2-0.5 µL of virus was injected into each hemisphere depending on the viral titration. Mice were allowed three weeks for recovery in their home cage before fitting with Neurologger 2A devices (see section below) and performing behavioral experiments. For experiments where temperature recordings were necessary, temperature loggers were usually inserted (abdominally) two to three weeks after mice had had their viral injection surgeries.

### EEG and EMG recordings and vigilance states scoring

Non-tethered EEG and EMG recordings were captured using Neurologger 2A devices (76). Screw electrodes were chronically inserted into the skull of mice to measure cortical EEG using the following coordinates: −1.5 mm Bregma, + 1.5 mm midline - first recording electrode; + 1.5 mm Bregma, −1.5 mm midline – second recording electrode; −1 mm Lambda, 0 mm midline – reference electrode). EMG signals were recorded by a pair of stainless steel electrodes implanted in the dorsal neck muscle. Four data channels (2 of EEG and 2 of EMG) were recorded with four times oversampling at a sampling rate of 200 Hz. The dataset was downloaded and waveforms visualized using Spike2 software (Cambridge Electronic Design, Cambridge, UK) or MATLAB (MathWorks, Cambridge, UK). The EEG signals were high-pass filtered (0.5 Hz,-3dB) using a digital filter and the EMG was band-pass filtered between 5-45 Hz (−3dB). Power in the delta (0-4 Hz), theta (6-10 Hz) bands and theta to delta band ratio were calculated, along with the root mean square (RMS) value of the EMG signal (averaged over a bin size of 5 s). All of these data were used to define the vigilance states of WAKE, NREM and REM by an automatic script. Each vigilance state was screened and confirmed manually afterwards. The peak frequency during NREM epochs were analyzed using Fourier transform power spectra to average power spectra over blocks of time.

### Core body temperature recordings

Core body temperature was recorded using temperature loggers (DST nano, Star-Oddi, HerfØlge, Denmark) implanted abdominally. A pre-defined program was set to sample the temperature data every two minutes for baseline core body temperature and drug/vehicle administration. At the end of the experiments, the loggers were retrieved and the data were downloaded and analyzed.

### Sleep deprivation and recovery sleep

The sleep deprivation protocol was similar to the one we used before (26), and it started at Zeitgeber time (ZT) zero (17:00), the start of the “lights-on” period when the sleep drive of the mice is at its maximum. Both experimental and control groups were sleep deprived for 5 hours by introducing novel objects or gently tapping on the cages. After sleep deprivation, mice were allowed to return back to their home cages for recovery NREM sleep. EEG and EMG signals together with temperature data were recorded for analysis.

### Chemogenetics and behavioral assessment

For chemogenetic activation, clozapine-N-oxide (CNO) (C0832, Sigma-Aldrich) was used. 1 mg/kg of CNO dissolved in saline or saline in same volume was administrated by intraperitoneal injection (*i.p.*) and the vigilance states and core body temperature were recorded. Mice were split into random groups that either received CNO or saline injection for an unambiguous comparison. Drugs were administrated at ZT18 (11:00, “lights-off”) when the mice were in their most active period and had their highest body temperature.

### Dexmedetomidine experiments

Prior to dexmedetomidine injection, animals with implanted temperature loggers were fitted with Neurologger 2A devices, and one hour of both baseline vigilance states and core body temperature was recorded as reference. 50 µg per kg of dexmedetomidine (Tocris Bioscience) was dissolved in saline and delivered *i.p.* at ZT19 (12:00, “lights-off”). Animals were placed back to their home cage immediately after injection for a further five-hour recording and the EEG, EMG and core body temperature were simultaneously recorded. A six-hour baseline recording from the same mouse of its natural sleep-wake cycle and core body temperature between ZT18 to ZT24 (11:00 −17:00, “lights-off”) was used for parallel comparison with the dexmedetomidine injection experiments.

### Immunohistochemistry

Mice were fixed by transcardial perfusion with 4% paraformaldehyde (Thermo scientific) in PBS, pH 7.4 after deep anesthesia by pentobarbital (100 mg/kg body weight; *i.p.*). Brains were removed and preserved in 30% sucrose in PBS. 35 or 50-µm-thick coronal were sliced using a Leica VT1000S vibratome. Free-floating sections were washed three times in PBS each for 5 minutes, permeabilized in 0.4% Triton X-100 in PBS for 30 minutes, blocked by incubation with 5% normal goat serum (NGS) (Vector) plus 0.2% Triton X-100 in PBS for 1 hour (all performed in room temperature) and then incubated with a GFP antiserum (rabbit, 1:1000, Life Technology, #A6455) or an mCherry antiserum (mouse, 1:1000, Clontech, #632543). Primary antisera were diluted in PBS with 2% NGS overnight at 4°C. The following day, primary antisera incubated sections were washed three times in PBS each for 10 minutes and subsequently incubated for 2 hours at room temperature in PBS with 2% NGS plus a dilution of an Alexa Fluor 488 goat anti-rabbit IgG (H+L) (1:1000, Molecular Probes, #A11034) or Alexa Fluor 594 goat anti-mouse IgG (H+L) (1:1000, Molecular Probes, #A11005). Sections were washed 4 times in PBS each for 10 minutes at room temperature and subsequently mounted on glass slides in Vectashield with DAPI (H-1200, Vector Laboratories).

### Quantification and statistical analysis

The sample sizes and statistical test for each experiment are stated in the figure legends. Origin v8.6 and Prism6 were used for statistical analyses.

## ACKNOWLEDGEMENTS

Our work was supported by the Wellcome Trust (107839/Z/15/Z, N.P.F. and 107841/Z/15/Z, W.W); the UK Dementia Research Institute (WW and NPF), the China Scholarship Council (YM), an Imperial College Schrödinger Scholarship (G.M.) and a studentship from the UK Medical Research Council (A.M.). The Facility for Imaging by Light Microscopy (FILM) at Imperial College London is in part supported by funding from the Wellcome Trust (grant 104931/Z/14/Z) and BBSRC (grant BB/L015129/1).

---

**Fig. S1.**
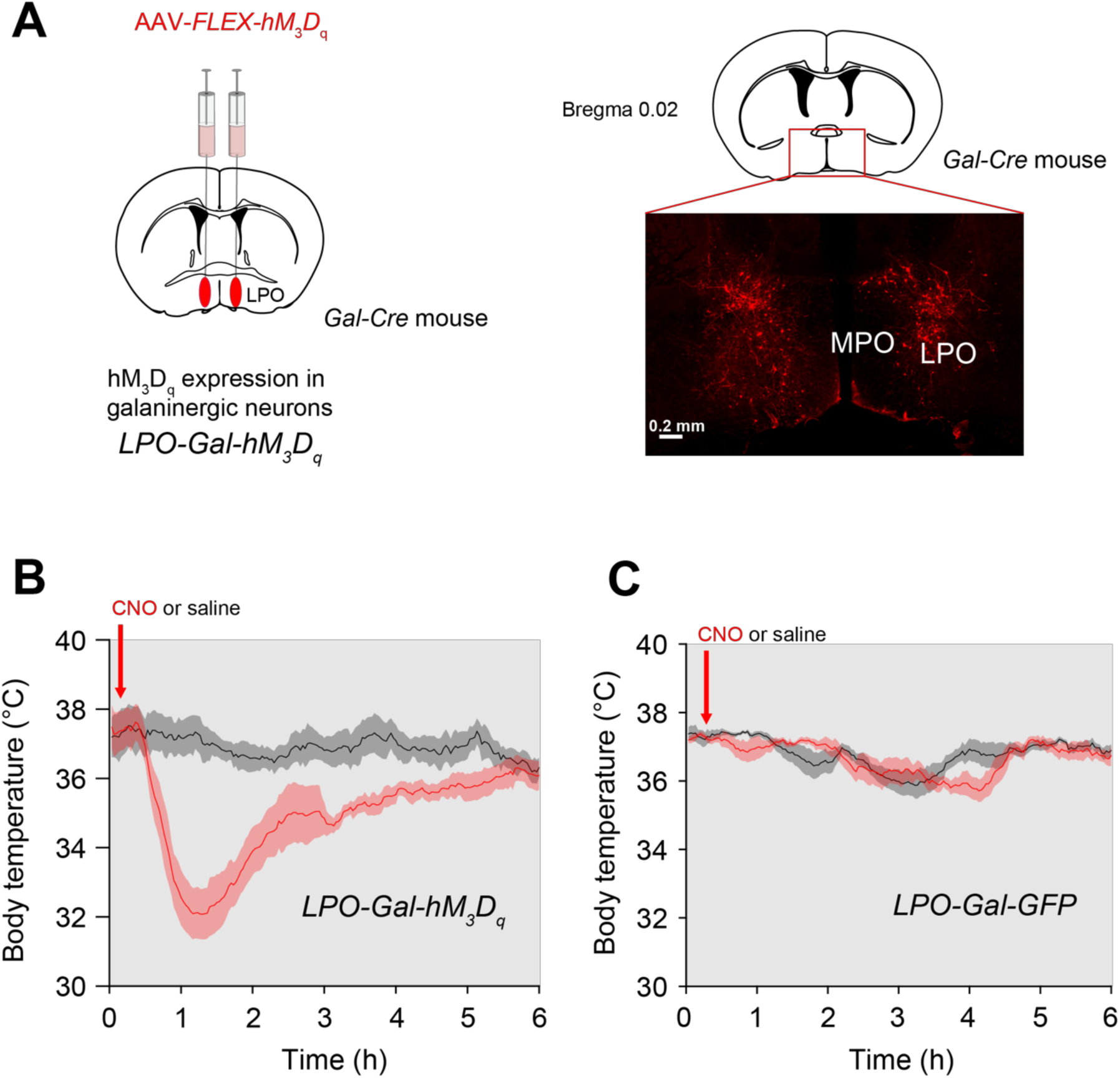
Selective chemogenetic activation of LPO^Gal^ neurons induces hypothermia. (A) Galanin neurons in the LPO of *Gal-Cre* mice were made selectively sensitive to CNO by bilaterally injecting AAV-*FLEX-hM_3_D_q_* to generate *LPO-Gal-hM_3_D_q_* mice. The hM_3_D_q_ receptor is fused to the mCherry protein, permitting visualization of receptor expression by immunohistochemistry with mCherry antibodies (image on right). Receptor expression was largely confined to the LPO area. (B) CNO (1 mg/kg) but not saline injection (*i.p.*) induced a strong acute hypothermia in *LPO-Gal-hM_3_D_q_* mice lasting several hours (*n*=5). (C) Control for off-target effects. CNO (1 mg/kg) injected (*i.p.*) into *Gal-Cre* mice (that had not been injected with AAV-*FLEX-hM_3_D_q_*) did not induce hypothermia compared with *Gal-Cre* mice that had received saline injections (*n*=5). All error bars represent the SEM.

**Fig. S2.**
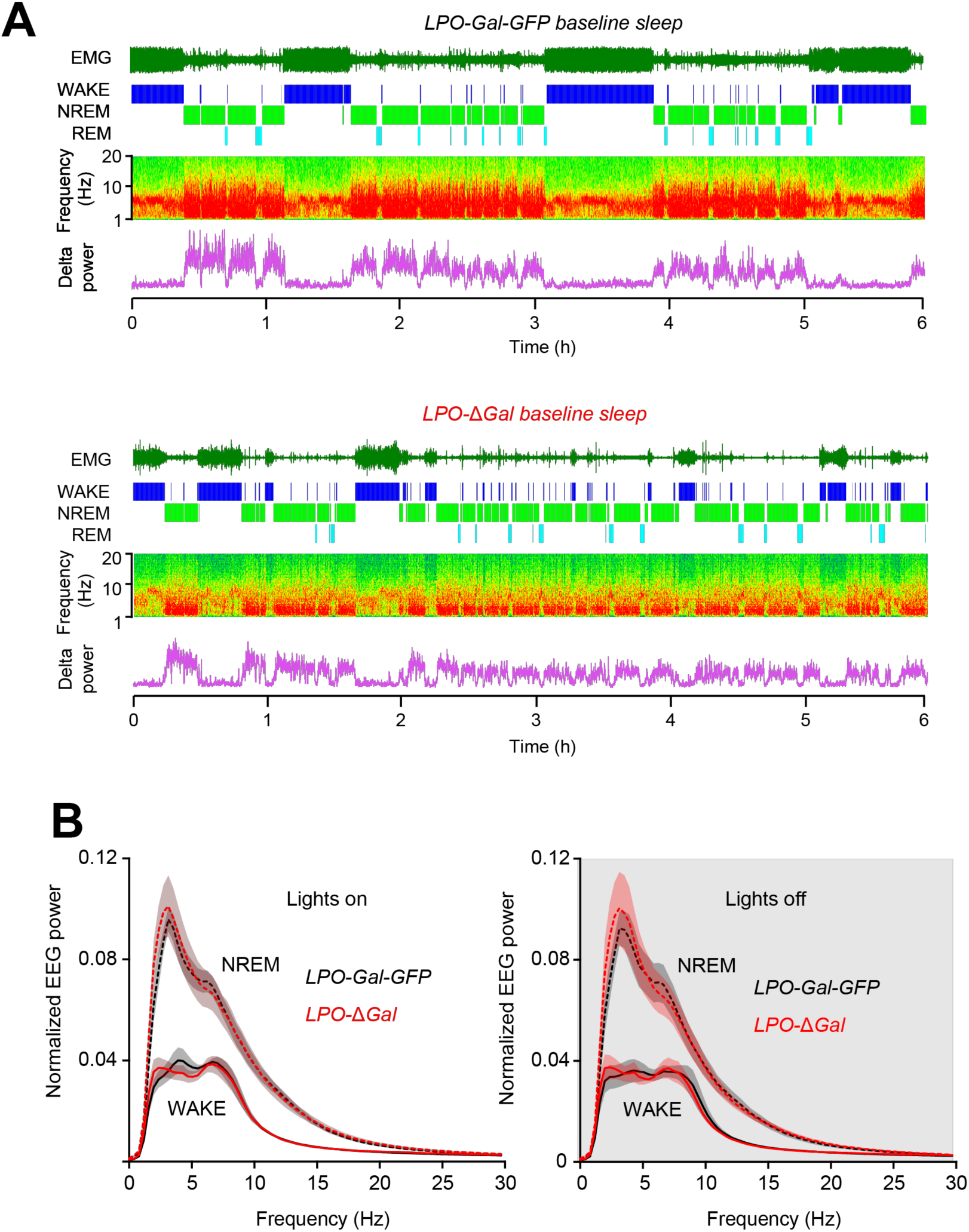
Baseline sleep in *LPO-Gal-GFP* mice and *LPO*-Δ*Gal* mice. (*A*) Examples of baseline EEG and EMG raw data and vigilance-state scoring for *LPO-Gal-GFP* mice and *LPO*-Δ*Gal* mice (from ZT0 to ZT6). (*B*) EEG power normalized such that the area under the curve was unity during the waking state. There were no significant differences in the baseline EEG power in either the WAKE state or NREM state between *LPO-Gal-GFP* mice and *LPO*-Δ*Gal* mice (*LPO-Gal-GFP*; *n*=8. *LPO-Δ*Gal; *n*=6). All error bars represent the SEM.

**Fig. S3.**
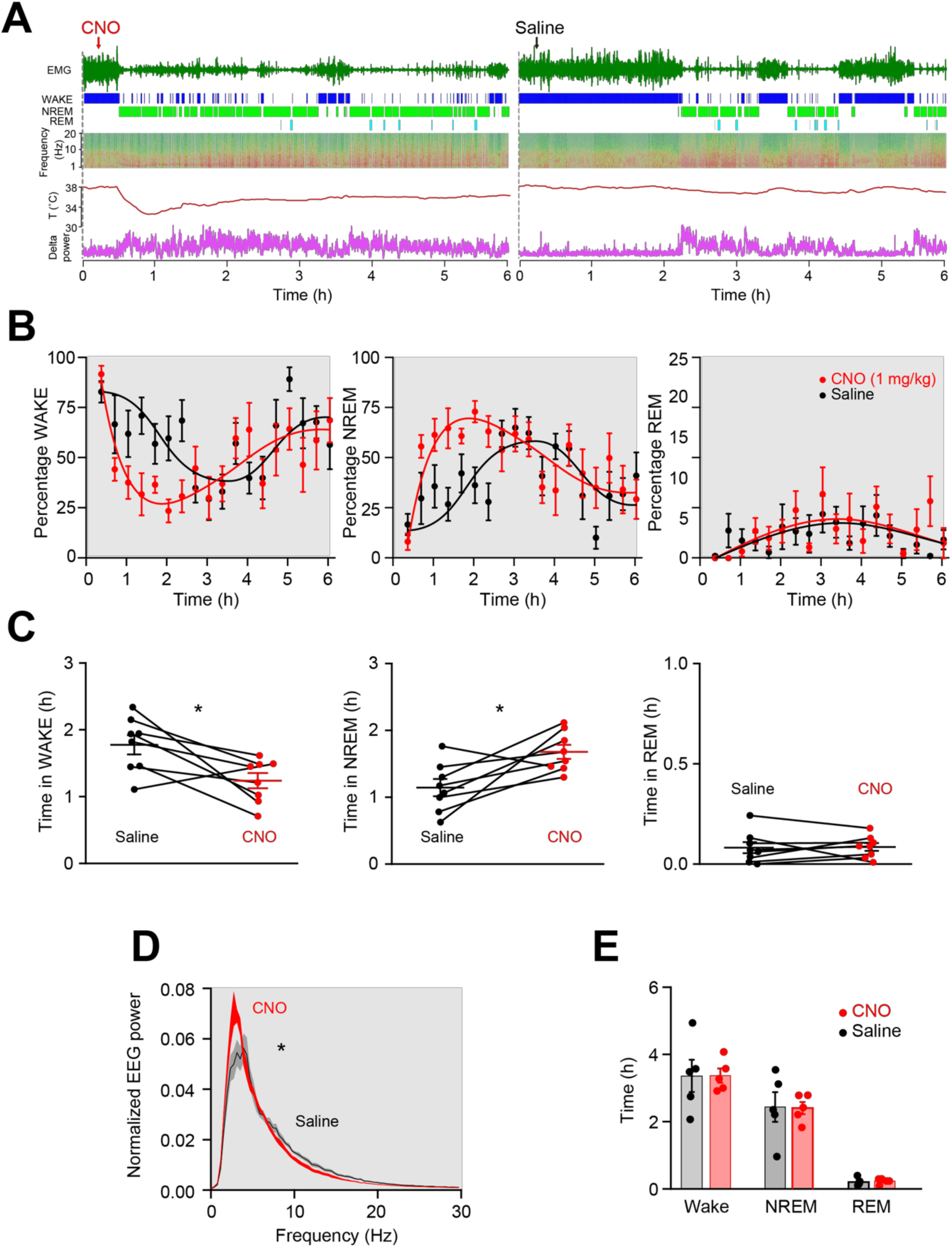
Selective chemogenetic activation of LPO galanin neurons by CNO increased NREM sleep. (*A*) Examples of EEG and EMG raw data and vigilance-state scoring, core body temperature and delta power for *LPO-Gal-hM_3_D_q_* mice following CNO (1 mg/kg) (left) or saline (right) *i.p.* injection. (*B*) The percentage of WAKE reduced and the percentage of NREM increased following CNO injection compared with control saline injection. The percentage of time in REM did not change (*n*=8). (*C*) Total time in WAKE, NREM and REM over the three hours following injection. (*n*=8, \**P*<0.05, paired two-tailed *t*-test) (*D*) The delta power of the CNO-induced NREM-like sleep had a significantly higher power than that of NREM sleep after saline injection (at the same time of injection) (\**P*<0.05, paired two-tailed *t*-test). Traces over the first 3 hours post injection. (*E*) Control for off-target effects: CNO (1 mg/kg) injected *i.p.* into *Gal-Cre* mice (that had not been injected with AAV-*FLEX-hM_3_D_q_*) did not induce NREM sleep above baseline amounts over the next 6 hours (compared with *Gal-Cre* mice that had received saline injections). All error bars represent the SEM.

